# Losing genes, gaining edits: how relaxed selection and inverted repeat expansion shape RNA editing in Schizaeaceae plastomes

**DOI:** 10.1101/2025.10.29.685334

**Authors:** Blake D. Fauskee, Li-Yaung Kuo, Farley M. Kwok van der Giezen, Kathleen M. Pryer

**Author notes:** These authors contributed equally to the study (Blake D. Fauskee, Li-Yaung Kuo).

## Abstract

RNA editing is a post-transcriptional pyrimidine exchange process that alters plastid and mitochondrial transcripts in nearly all land plants. Although confined to organelles, it is directed by nuclear-encoded PLS-type pentatricopeptide repeat (PPR) proteins, each typically recognizing a specific RNA target. While many editing sites are functionally neutral, edits at cryptic start and internal stop codons have been implicated in modulating organellar gene expression. Ferns—and some lycophytes—are unique among vascular plants in exhibiting both C-to-U and U-to-C editing, making them valuable for studying the evolution of both forms. Here, we examine chloroplast RNA editing in four Schizaeales species (*Schizaea dichotoma, Actinostachys digitata, Anemia phyllitidis, Lygodium microphyllum*), two of which possess non-photosynthetic gametophytes, providing a natural contrast with fully photosynthetic relatives. Despite extensive plastome reduction, including loss of the *ndh* suite and, in *Actinostachys*, all *chl* genes, *Schizaea* and *Actinostachys* exhibit dramatically elevated numbers of C-to-U edits. Genes evolving under relaxed selection accumulate more editing sites, and editing abundance per gene correlates with the magnitude of relaxed constraint, suggesting relaxed selection promotes edit proliferation. *S. dichotoma* and *A. digitata* also show expansion of the chloroplast inverted repeat (IR), and genes translocated into the IR exhibit reduced substitution rates and higher editing densities, indicating that IR expansion slows the loss of edits. Finally, annotation of PPR proteins revealed few full-length editing factors, consistent with catalytic domains assembling *in trans* and highlighting the modular nature of the fern editosome.

## Introduction

Nearly all land plants undergo RNA editing, a post-transcriptional process that introduces targeted nucleotide modifications to RNA transcripts derived from organellar genomes. This editing process yields mRNA sequences that no longer match their DNA templates (reviewed in Knoop 2023). Most RNA editing events in organellar genomes act to restore evolutionarily conserved amino acid sequences to faithfully yield functional proteins (reviewed in Small et al. 2020). Two forms of RNA editing occur in land plant organelles: C-to-U RNA editing, where cytidines are deaminated to yield uridines, and U-to-C editing, where uridines are converted to cytidines via a suspected transamination reaction (reviewed in Chateigner-Boutin and Small 2011; Ichinose and Sugita 2017; Hayes et al. 2024). C-to-U editing restores protein functionality in the organellar genomes of all land plants, except for the liverwort *Marchantia polymorpha* (Rüdinger et al. 2008; Shen et al. 2024). U-to-C editing presents a different evolutionary picture where it is absent in seed plants (Tillich et al. 2006), but retained in hornworts (Kugita et al. 2003), some lycophyte lineages (Grewe et al. 2011; Kwok van der Giezen et al. 2024), and ferns (Wolf et al. 2004; Guo et al. 2015; Fauskee et al. 2025).

Although RNA editing in plants occurs primarily in organellar genomes, it is carried out by nuclear-encoded pentatricopeptide repeat (PPR) proteins (Lurin et al. 2004; Chateigner-Boutin et al. 2008). PPR proteins are a large family of RNA-binding proteins that play diverse roles in organellar RNA metabolism, including editing, transcript stabilization (Prikryl et al. 2011), splicing (de Longevialle et al. 2007; Chateigner-Boutin et al. 2011), and endonucleolytic cleavage (Huynh et al. 2023). These proteins are composed of tandem arrays of helix-turn-helix units, known as PPR motifs, that form a super-helical structure with the central RNA binding groove. Individual PPR motifs typically recognize one RNA nucleotide at a time, with binding specificity conferred by hydrogen bonds formed between the 5th and C-terminal amino acid residues in each PPR motif and the corresponding RNA base (Barkan et al. 2012). PPR proteins are present throughout eukaryotes, but the gene family has undergone dramatic expansion in land plants, where these proteins have been co-opted for highly specialized roles in organellar gene expression (Gutmann et al. 2020). Land plant PPR proteins are broadly divided into two classes based on their motif architecture and function. P-class PPR proteins consist of tandem repeats of a 35-amino-acid motif and primarily function in RNA stabilization, splicing, and cleavage. In contrast, PLS-class PPR proteins function as RNA editing enzymes and contain repeating triplet units of P1 (35 amino acids), L1 (35 amino acids), and S1 (31 amino acids) motifs, often accompanied by additional P2 (35 amnio acids), L2 (36 amino acids), and S2 (32 amino acids) variants. These proteins are typically capped by C-terminal catalytic domains— either E1-E2-DYW for C-to-U editing, or E1-E2-DYW:KP for U-to-C editing—that carry out the nucleotide base conversions necessary for producing functional transcripts.

Although some RNA editing sites––notably C-to-U edits that restore start codons and U-to-C edits that correct internal stop codons––are evolutionarily conserved for their adaptive roles in regulating plastid gene expression (Fauskee et al. 2025), most editing sites likely evolve through a neutral evolutionary ratchet mechanism known as constructive neutral evolution (CNE). This is where otherwise deleterious mutations in organellar DNA are tolerated and reach fixation because they are post-transcriptionally corrected through RNA editing (Covello and Gray 1993, Lukeš et al. 2011). Under the CNE model, new RNA editing sites proliferate when organellar genome mutations are compensated by pre-existing nuclear-encoded PLS-class PPR editing proteins, which post-transcriptionally restore protein function through editing. Despite the potential for RNA editing sites to accumulate neutrally, several comparative studies have shown progressive loss of editing sites in the chloroplast genomes of angiosperms (Mower 2008; Ishibashi et al. 2019) and several fern lineages (Fauskee et al. 2025). This trend likely reflects mutational bias in AT-rich chloroplast genomes, which favors loss of C-to-U editing sites, the only editing type in angiosperms and the predominant type in ferns. C-to-T transitions occur at a particularly high rate in chloroplast genomes (Huang et al. 2005). When these mutations arise at C-to-U editing sites, they restore the ancestral codon sequence, thereby eliminating the need for editing at that position. Thus, while a small subset of RNA editing sites are conserved by selection, the overall evolutionary trend of RNA editing in most plant lineages is one of neutral gain outpaced by progressive loss.

Despite recent advances in our understanding of plastid RNA editing (Wolf et al. 2004; Guo et al. 2015; Ishibashi et al. 2019; Gerke et al. 2020; Small et al. 2020; Fauskee et al. 2025), little is known about how this process operates in plants exhibiting partial or complete heterotrophy during their life cycle. Likewise most research on plastome evolution has centered on green plants with highly conserved plastome architecture (Mower and Vickrey 2018). With few exceptions, land plant plastomes are remarkably stable in size, structure, and gene content: most are circular-mapping molecules (Rochaix 1978) ranging from 120– 160 kb in length and exhibit a quadripartite structure consisting of a large and small single-copy region (LSC and SSC, respectively) separated by two inverted repeats (IR_A_ and IR_B_) (Ruhlman and Jansen 2014; Mower and Vickrey 2018). Plastomes of fully autotrophic plants typically encode approximately 80 protein-coding genes (Wicke et al. 2011), four rRNAs (consistently located in the IR; Zhu et al. 2016), and around 30 tRNAs (Jansen and Ruhlman 2012). In contrast, plastomes of non-photosynthetic plants—where both the sporophyte and gametophyte lack photosynthetic ability—and partially non-photosynthetic plants—where only one life stage retains photosynthesis—frequently exhibit significant departures from this conserved genomic organization. Research on independently derived parasitic and mycoheterotrophic lineages has revealed extensive and rapid gene loss (particularly *ndh* genes), genome size reduction, and structural rearrangements, primarily driven by relaxed purifying selection on genes associated with photosynthesis (Barrett et al. 2014; Bellot and Renner 2016; Wicke et al. 2016). These pronounced differences raise an important and largely unexplored question: how does plastid RNA editing evolve under conditions of plastome deterioration and diminished selective pressure on photosynthetic function?

To our knowledge, there is only one study to date that has investigated the plastid editome of a non-photosynthetic plant. Funk et al. (2007) identified plastid RNA editing sites in two parasitic species of *Cuscuta—C. reflexa* and *C. gronovii*—revealing reduced editing levels in both relative to their close photosynthetic relatives. While *Cuscuta* represents an important case of plastome degradation and functional gene loss associated with a parasitic lifestyle, the utility of angiosperms for broader evolutionary studies of plastid RNA editing is limited in that they exclusively possess C-to-U editing and retain relatively few editing sites—typically 30 to 50 per plastome (reviewed in Small et al. 2020). This limits investigation of evolutionary patterns in the diversity, directionality, and abundance of plastid RNA editing.

Ferns and hornworts, by contrast, are the two most extensively studied plant lineages exhibiting abundant plastid RNA editing, with both C-to-U and U-to-C editing types present (Kugita et al. 2003; Guo et al. 2015; Villareal et al. 2018; Fauskee et al. 2025). Ferns are unique among land plants in that they have free-living sporophyte and gametophyte generations each with distinct physiological roles (reviewed in Krieg and Chambers 2022). In almost all fern lineages, both life stages are photosynthetically active. The Schizaeaceae (order Schizaeales) are exceptional; several groups produce achlorophyllous gametophytes, while members of the other families in the order—Anemiaceae and Lygodiaceae—retain green gametophytes (Bierhorst 1975). The plastomes of Schizaeaceae exhibit strikingly similar genomic reductions and structural rearrangements to those seen in heterotrophic angiosperms: all members have lost the entire suite of plastid-encoded NAD(P)H dehydrogenase complex (*ndh*) genes, while *Actinostachys* has additionally lost the light-independent protochlorophyllide reductase (*chl*) genes (Labiak and Karol 2017; Ke et al. 2022). This well-characterized variation in gametophyte plastid function and plastome structure in Schizaeales provides a powerful comparative framework for examining how both C-to-U and U-to-C RNA editing evolves under conditions that differ markedly from fully photosynthetic plants—specifically in lineages where plastid activity is developmentally partitioned and the plastomes themselves have undergone extensive structural changes.

Here, we report on the plastid editomes of four species in the order Schizaeales: two with achlorophyllous gametophytes—*Schizaea dichotoma* (L.) Sm. and *Actinostachys digitata* (L.) Wall. (Schizaeaceae)—and two with green, photosynthetic gametophytes—*Anemia phyllitidis* (L.) Sw. (Anemiaceae) and *Lygodium microphyllum* (Cav.) R. Br. (Lygodiaceae). We present newly assembled and annotated plastomes, validate and characterize plastid RNA editing sites using transcriptome data, assess the abundance of PPR proteins, and apply phylogenetic and bioinformatic approaches to investigate how both C-to-U and U-to-C RNA editing evolves in relation to plastome structure and gametophyte photosynthetic capacity.

## Methods

### Sampling, DNA extraction and sequencing, plastome assembly and annotation

DNA was extracted from four species in three families in the order Schizaeales: *Schizaea dichotoma* and *Actinostachys digitata* (Schizaeaceae), *Anemia phyllitidis* (Anemiaceae), and *Lygodium microphyllum* (Lygodiaceae). DNA isolations were performed using the E.Z.N.A. SP Plant and Fungal DNA Extraction Kit from Omega Bio-Tek (Omega Bio-Tek, Norcross, GA, USA; D5511). Isolated DNA was then quantified using a Qubit 2 Fluorometer (Thermo Fisher Scientific Inc., Walden, MA, USA) and the Qubit dsDNA High Sensitivity Quantification Assay kit (Thermo Fisher Scientific Inc., Walden, MA, USA; Q32851). DNA libraries for Illumina sequencing were then constructed using the NEBNext Ultra II DNA Library Prep Kit for Illumina (New England Biosciences, Ipswitch, MA, USA; E7645) following the manufacturer’s default protocol for 200 base-pair inserts. To enable multiplexing, each sample was tagged with a unique barcode using the NEBNext Multiplex Oligos for Illumina (New England Biosciences, Ipswitch, MA, USA; E6609). Resulting DNA libraries were sequenced on the Illumina Novaseq X Plus by Novogene Co., Ltd (Beijing, China) using 150 base-pair, paired-end chemistry. De-multiplexing was also performed by Novogene Co., Ltd.

Paired-end DNA reads were uploaded to the Duke Compute Cluster (Duke University, Durham, NC, USA) where adapters were trimmed and low-quality reads were removed using Trimmomatic version 0.39 (Bolger et al. 2014) with the following settings: LEADING:3 TRAILING:3 SLIDINGWINDOW:4:15 MINLEN:36. Using the resulting trimmed and paired reads, chloroplast genomes were assembled *de novo* using NOVOPlasty version 4.3.3 (Dierckxsens et al. 2017). For each assembly, an *rbcL* sequence was obtained from perviously published full plastomes uploaded in GenBank for the same species (but different voucher) and used as the seed sequence. Specifically, ON207052 for *Schizaea dichotoma*, ON207049 for *Actinostachys digitata*, OM990738 for *Anemia phyllitidis*, and MG761729 for *Lygodium microphyllum*. Draft plastomes were then polished iteratively using Pilon (Walker et al. 2014) until Pilon suggested no further changes to the assembly. Polished plastome assemblies were then annotated in Geneious version 2022.0.2 using the BLAST plugin followed by manual adjustment. The same Genbank vouchers from which seed sequences were obtained were used as the annotation reference.

### Transcriptome sequencing and assembly

RNA was extracted from the same vouchers for which plastome assemblies were generated. The RNA was extracted from either fresh or flash-frozen green sporophyte tissue. The RNA extractions were performed using the E.Z.N.A. Plant RNA Kit from Omega Bio-Tek (Omega Bio-Tek, Norcross, GA, USA; R6827). During RNA extraction, samples were additionally treated with DNase I to reduce DNA contamination. DNase I treatments were performed with the Millipore Sigma DNase I kit (Millipore Sigma, Darmstadt, Germany; 69182). RNA extractions were quantified using the Qubit 2 Flurometer (Thermo Fisher Scientific Inc., Walden, MA, USA) and the Qubit RNA High Sensitivity Quantification Assay kit (Thermo Fisher Inc., Walden, MA, USA; Q32852).

RNA (cDNA) libraries were constructed using the Zymo-Seq RiboFree Total RNA Library Kit (Zymo Research, Irvine, CA, USA; R3000) or the NEBNext Ultra II Library Prep (New England Biosciences, Ipswich, MA, USA; E7775), with ribosomal depletion probes designed for plant samples supplied by New England Biosciences as part of a beta test agreement. For both kits, library preparation was performed following suggested manufacturer’s protocols. Unique barcodes were ligated to each sample to enable multiplexing, and the resulting libraries were sequenced by Novogene Co., Ltd (Beijing, China) using 150 base-pair, paired-end chemistry on the Illumina NovaSeq X Plus. All data quality control and demultiplexing was carried out by Novogene. Raw transcriptomic and genomic data generated for this project are available from the Sequence Read Archive (SRA), BioProject number: XXX

### De novo transcriptome assembly

Raw transcriptomic data was trimmed of adapters using BBduk from the BBmap suite (Bushnell 2014) with settings (ktm=r k=23 mink=11 hdist=1 ftm=5 tpe tbo). Trimmed reads were assembled using rnaSPAdes v4.0 with in assembler-only mode (Bushmanova et al. 2019). The completeness of the resulting transcriptome assemblies was assessed with Benchmarking using single copy orthologues (BUSCO), referencing the embryophyta_odb10 dataset (Simão et al. 2015).

### Identification of pentatricopeptide repeat protein sequences

Transcriptomes for each species were scanned for open reading frames (ORFs) in all six (three forward, three reverse) reading frames using the Julia script orfinder.jl (Kwok van der Giezen et al., 2025) and translated to a protein FASTA file. Models of PPR proteins were identified in the ORFs using ‘hmmsearch’ (HMMER v3.2.1) and the “DYW/DYW:KP” hidden Markov model for PPR proteins described in (Gutmann et al. 2020). PPRFinder (Gutmann et al. 2020) was run on the hmmsearch output to generate files containing annotated PPR motif and domain structures.

### RNA editing detection

To detect plastid RNA editing sites in the four Schizaeales plastomes, all protein coding gene sequences for each species were extracted using Geneious (Kearse et al. 2012) along with 100 base-pairs flanking the beginning and end of each gene. For each species, protein coding genes along with flanking regions were combined into a single multi-fasta file. A custom RNA editing detection pipeline modified from Edera and Sanchez-Puerta (2021) was implemented to detect potential RNA editing sites. In this pipeline, RNA reads are trimmed twice with Trimmomatic (version 0.39), first in paired-end mode using the following settings: LEADING:3 TRAILING:3 SLIDINGWINDOW:4:15 MINLEN:36 to remove adapters and low-quality reads, then again in single-end mode using the HEADCROP:13 setting which removes the first 13 bases in each read where GC content was non-uniform. RNA reads were then mapped to the plastid protein-coding genes for each species using Bowtie2 version 2.2.4 (Langmead and Salzberg 2012). A BAM file was then generated using Samtools version 1.14 (Li et al. 2009). The total number of RNA reads mapped to each site in each gene, as well as the number of reads with each nucleotide present at that site, were calculated using bam-readcount version 0.8.0 (Khanna et al. 2022) and custom Linux commands. This workflow ultimately output a TSV file showing the base present in the DNA at every site in every gene analyzed, the total number of mapped RNA reads to each site, and the number of mapped reads displaying each of the four nucleotides. An example script for these steps is available on GitHub (https://github.com/bfauskee/fauskee-fern-rna-editing-scripts).

Using the R pipeline described in Fauskee et al. (2025), putative RNA editing sites were detected and characterized. An example of this R pipeline is available on Github (https://github.com/bfauskee/fauskee-fern-rna-editing-scripts). Sites were considered RNA editing sites if they met the following conditions: at least 10 RNA reads mapped to the site and at least three reads and 10% of the total mapped RNA reads had the edited base (for example, a T mapped to a C for a C-to-U editing site). Uncommon regions with very low RNA read coverage were manually inspected. This pipeline also characterizes the amino acid change induced by each RNA editing event, the codon position it occurs at, the efficiency of each editing site, along with several other features. RNA editing efficiency is defined as the proportion of mapped RNA reads displaying the edited base (e.g., the number of reads with a T mapped to a C divided by the total number of reads mapped to that site).

### Evolutionary analyses of RNA editing sites

To understand how RNA editing sites are evolving in the Schizaeales, we performed several evolutionary and phylogenetic analyses. First, we aligned the coding regions of all plastid protein-coding genes for each of the four Schizaeales species using MAFFT version 7.505 (Katoh and Standley 2013). We then used the alignment position of each site to compare the presence and absence of each unique RNA editing site across the four species by generating UpSet plots using the R package UpSetR version 1.4 (Conway et al 2017).

Next, we aligned the coding regions of each plastid protein-coding gene again using MAFFT version 7.505 (Katoh and Standley 2013), but with the addition of the previously published plastome of *Salvinia cucullata* (Genbank accession number: MF177095.1). Genes were concatenated and partitioned by gene using AMAS (Boroweic 2016) to generate a supermatrix. We then carried out maximum likelihood (ML) tree estimation on the concatenated plastid supermatrix in IQ-TREE2 version 2.2.2.7 (Minh et al. 2020) using 1,000 ultrafast bootstrap replicates. Substitution model selection and optimal partitioning schemes were automated in IQ-TREE2 using the MFP+MERGE option. For substitution model selection, the model preferred by the Bayesian information criterion (BIC) for each partition model was used. The concatenated ML tree was then rooted with *S. cucullata*, which was then pruned from the tree using the Phyx package version 1.3 (Brown et al. 2017).

We also calculated nucleotide diversity (π) to test whether differences in RNA editing among species reflect off-target edits. Using the Schizaeales-only multiple sequence alignments described previously, we created additional “post-edited” alignments by recoding RNA editing sites as the post-edited base (changing a C to a T for a C-to-U edit, and a T to a C for a U-to-C edit) without re-aligning the sequences. These “post-edited” alignments represent the mRNA sequences. We then calculated pairwise π for all genes across all six species combinations on both the unedited DNA alignments and the recoded RNA alignments using custom R scripts. Lastly, we plotted DNA π versus RNA π.

To assess whether selection is relaxed in any plastid genes in Schizaeaceae relative to Anemiaceae and Lygodiaceae, we expanded our dataset to include several additional Schizaeales plastomes from Genbank. These included: *Schizaea elegans* (KX258660), *Schizaea pectinata* (NC035808), *Actinostachys pennula* (KU764518), *Lygodium flexuosum* (OM350009), *Lygodium merrillii* (OM350010), *Lygodium japonicum* (KC356645), and *Lygodium circinatum* (OM327797). Using the same alignment, concatenation, and phylogenetic inference methods outlined above, we constructed another phylogenetic tree for our expanded dataset. This resulting ML tree was similarly rooted with *S. cucullata* and then *S. cucullata* was pruned. We then assessed whether selection was relaxed in any plastid protein-coding genes in Schizaeaceae relative to Anemiaceae and Lygodiaceae. Here only genes present across all species were analyzed, resulting in 69 protein-coding genes. The selection analysis was carried out using RELAX (Wertheim et al. 2015) implemented in HYPHY version 2.5.64 (Kosakovsky Pond et al. 2020). For genes with relaxed selection (k value significantly less than 1), we compared the k value (lower values representing more extreme relaxation of selection) to the pairwise difference in the number of nonsynonymous C-to-U edits between each Schizaeaceae and non-Schizaeaceae species. This yielded four pairwise comparisons: *Schizaea dichotoma* vs *Anemia phyllitidis, S. dichotoma* vs *Lygodium microphyllum, Actinostachys digitata* vs *A. phyllitidis*, and *A. digitata* vs *L. microphyllum*. For each pairwise comparison we plotted the pairwise C-to-U editing count difference by the k value for genes that had a k value significantly less than 1 (p < 0.05). We performed a multiple linear regression to quantify the relationship between k and the pairwise editing difference with gene length as a covariate in R (R Core Team 2021).

We also investigated whether genes translocated into the inverted repeat in the Schizaeaceae exhibited decelerated substitution rates, as has been observed in other fern lineages (Li et al. 2016). We constructed individual gene trees for each chloroplast gene using our expanded sampling dataset outlined above. The gene trees were built in IQ-TREE2 (Minh et al. 2020) with substitution model selection carried out by using the MFP option. Again, the substitution model preferred by BIC was used. Each gene tree was then rooted with *S. cucullata* and then *S. cucullata* was pruned. For each gene tree, we calculated the mean relative distance to the most recent common ancestor (MRCA) for *Schizaea, Actinostachys*, and *Lygodium. Anemia* was excluded here because we only had access to one *Anemia* species. The mean relative distance to the MRCA for each genus was calculated by taking the mean branch length distance to the MRCA for each *Schizaea, Actinostachys*, or *Lygodium* species and dividing that mean by the total tree length. This enabled meaningful comparisons across genes. Distances were extracted from trees using the distRoot function in the adephylo R package version 1.16 (Jombart et al. 2010). We then binned genes by whether they were found consistently in single copy regions (small or large single copy), consistently found in the IR, found in the IR in both *Actinostachys* and *Schizaea*, or found in the IR only in *Schizaea*. We then plotted the mean relative MRCA distance for each of the three genera for each gene.

## Results

### Plastome structure and content in Schizaeales

Our plastome assemblies from four Schizaeales species revealed remarkable diversity in both size and gene content. The plastomes of *Anemia phyllitidis* and *Lygodium microphyllum* are similar in size, measuring 163,673 bp and 163,286 bp, respectively (Fig. 1). By contrast, the plastomes of the two Schizaeaceae species are considerably smaller. *Schizaea dichotoma* has a plastome of 160,380 bp, while *Actinostachys digitata* has a notably reduced plastome at 135,863 bp (Fig. 1).

**Figure 1.**
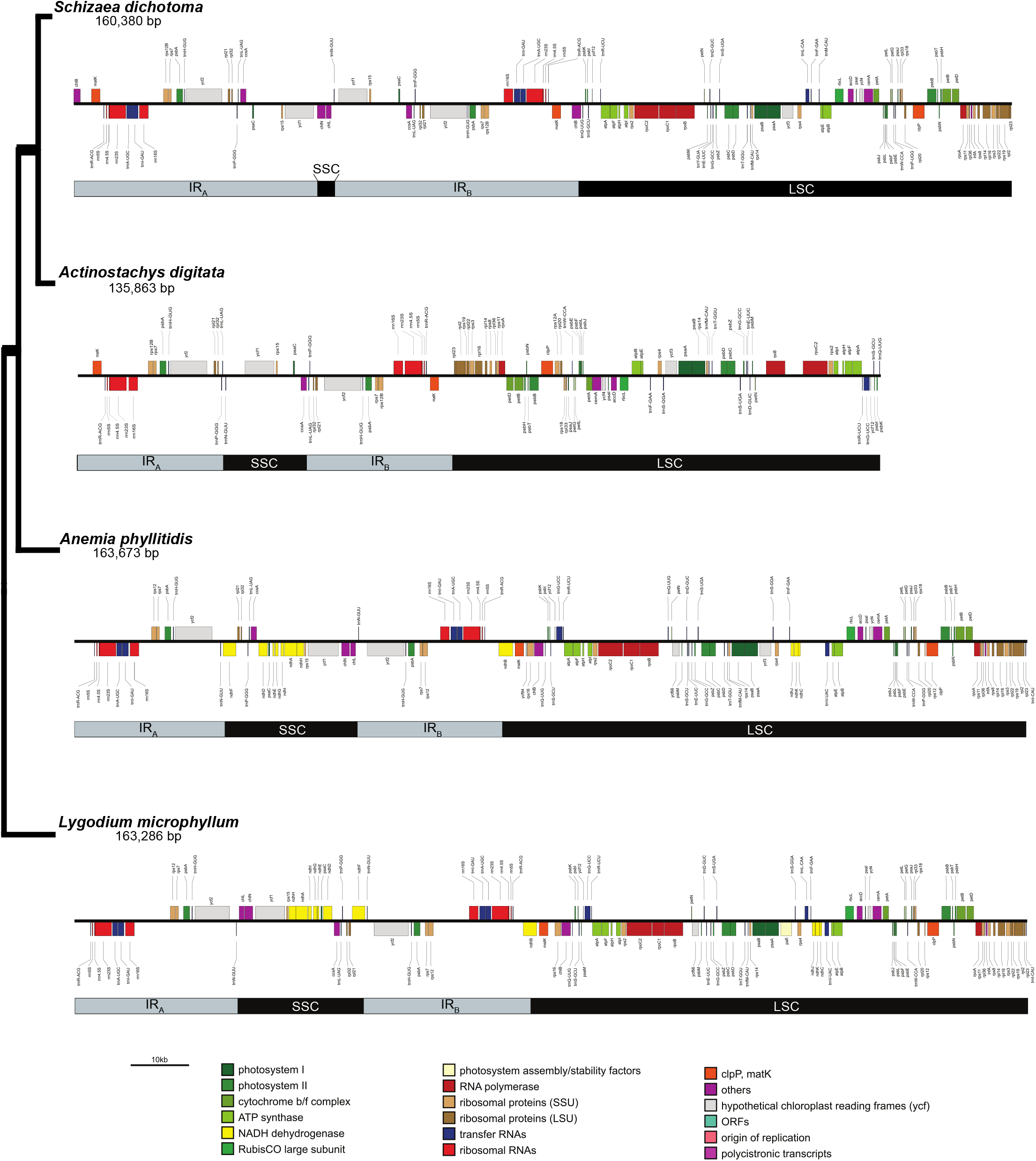
Linearized plastome maps for four Schizaeales species. Inverted repeat (IR) regions are outlined in grey and single copy regions (the SSC and LSC) in black. Phylogenetic relationships of the four species are show on the left.

In Schizaeaceae, we observed dramatic expansion of the inverted repeat (IR) regions accompanied by substantial reduction of the small single copy (SSC) region. *Lygodium microphyllum* and *A. phyllitidis* share a typical SSC structure, containing the same 14 protein-coding genes, a portion of *ndhF*, and two tRNAs (Fig. 1). In *A. digitata*, the IR has expanded slightly to incorporate *rps21* and *rps32*, which reside in the SSC in *L. microphyllum* and *A. phyllitidis* (Fig. 1). IR expansion in *S. dichotoma* is even more extreme: its SSC is reduced to just 2,863 bp and includes only *chlL, chlN*, and *trnN-GUU* (Fig. 1). This expansion brings *ccsA, psaC*, rps15, and *ycf1* into the IR—genes that remain in the SSC in the other three species (Fig. 1). Additionally, *matK* has been translocated into the IR in both Schizaeaceae species, while it remains in the large single copy (LSC) region in *L. microphyllum* and *A. phyllitidis* (Fig. 1).

Extensive plastid gene loss was observed in Schizaeaceae. Both *S. dichotoma* and *A. digitata* have lost the entire suite of 11 NAD(P)H dehydrogenase (*ndh*) genes for electron transport, as well as *rps16* and *ycf66* (Fig. 2). In addition, *A. digitata* has lost all three genes associated with chlorophyll biosynthesis: *chlB, chlF*, and *chlN* (Fig. 2). The Photosystem I *psaM* gene is absent from all plastomes assembled here except for *L. microphyllum* (Fig. 2). In total, we detected 86 protein-coding genes in *L. microphyllum*, 85 in *A. phyllitidis*, 72 in *S. dichotoma*, and 69 in *A. digitata* (Fig. 2). To determine whether the missing *ndh* genes might have been functionally transferred to the nucleus in Schizaeaceae, we searched the assembled transcriptomes of *A. digitata* and *S. dichotoma*, but found no evidence of these genes, suggesting that they have been entirely lostfrom Schizaceae. Likewise, the missing *chl* genes from *A. digitata*, were also entirely lost.

**Figure 2.**
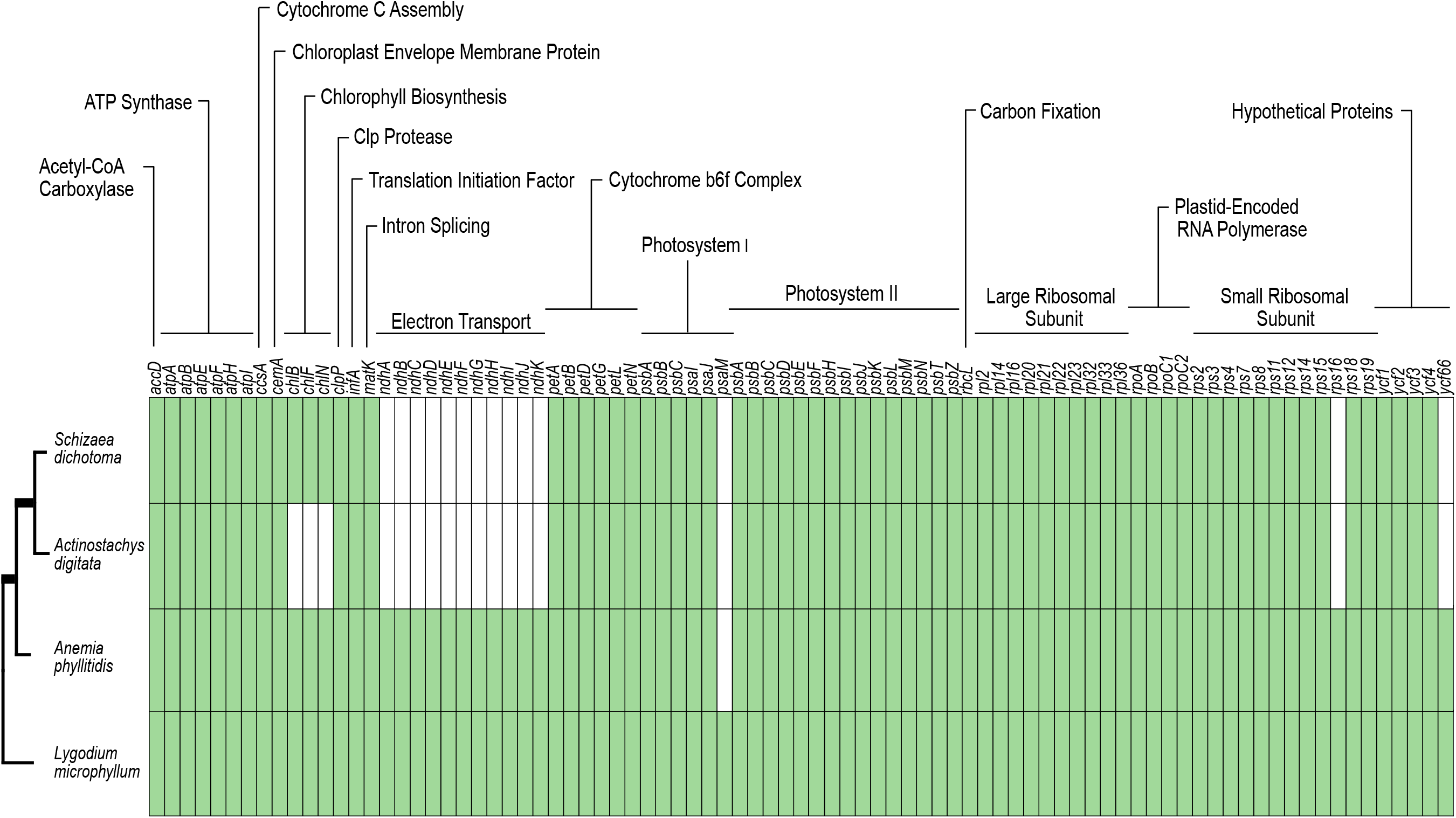
Gene presence/absence in four plastomes of Schizaeales. Phylogenetic relationships of the four species are show on the left. Green cells represent the presence of a given gene while white cells represent absence. General functional groups are shown above the gene names.

### Plastid RNA editing evolution

Analyses of chloroplast RNA editing in the Schizaeales revealed a sharp disparity between the two edit types. C-to-U editing levels were extremely variable ranging from as low as 286 C-to-U edits in *L. microphyllum* to 631 C-to-U edits in *S. dichotoma*, with both Schizaeaceae species displaying higher C-to-U editing levels than the non-Schizaeaceae species (Fig. 3). U-to-C editing levels, however, showed little variation ranging from 37 edits in *A. digitata* to 42 edits in *S. dichotoma* (Fig. 3). The number of edits per protein-coding base (CDS) in the plastomes was additionally calculated and mirrors the patterns of raw edit numbers in each lineage with the highest number of C-to-U edits per CDS in *S. dichotoma* and *A. digitata*, but little variation across the order for U-to-C edits (Fig, 3).

**Figure 3.**
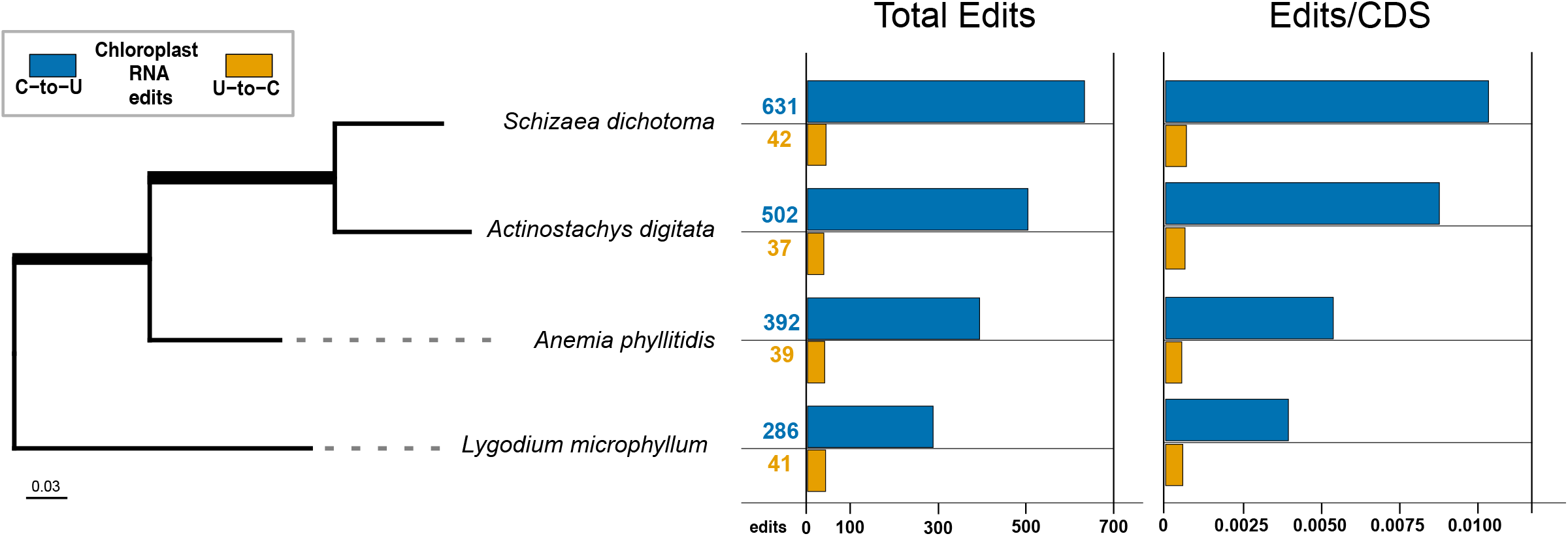
Chloroplast RNA editing levels in the Schizaeales. Phylogenetic relationships of the four species are shown on the left with branch lengths from a maximum likelihood analysis. Branch lengths are expressed in substitutions/site and a scale bar is in the bottom-left corner. Blue horizontal bars show the number of C-to-U edits in the leftmost panel under “Total Edits” and the number of C-to-U edits per the number of protein coding bases in the chloroplast genome in the rightmost panel under “Edits/CDS”. Similarly gold bars indicate the number of total U-to-C edits or U-to-C edits per coding base on the left and right, respectively.

Most C-to-U edits in all four species resulted in nonsynonymous codon changes (Fig. 4). A smaller proportion restored start codons by converting ACG to AUG, while others are synonymous and did not alter the amino acid sequence (Fig. 4). By contrast, most U-to-C edits restored coding potential by converting internal stop codons to a sense codons, while a few resulted in nonsynonymous amino acid substitutions (Fig. 4). Like C-to-U edits, a small proportion are synonymous (Fig. 4). Nonsynonymous and synonymous U-to-C edits tend to be evenly distributed across transcripts while U-to-C edits that correct internal stop codons tend to be biased toward the 5’ end (Fig. 4).

**Figure 4.**
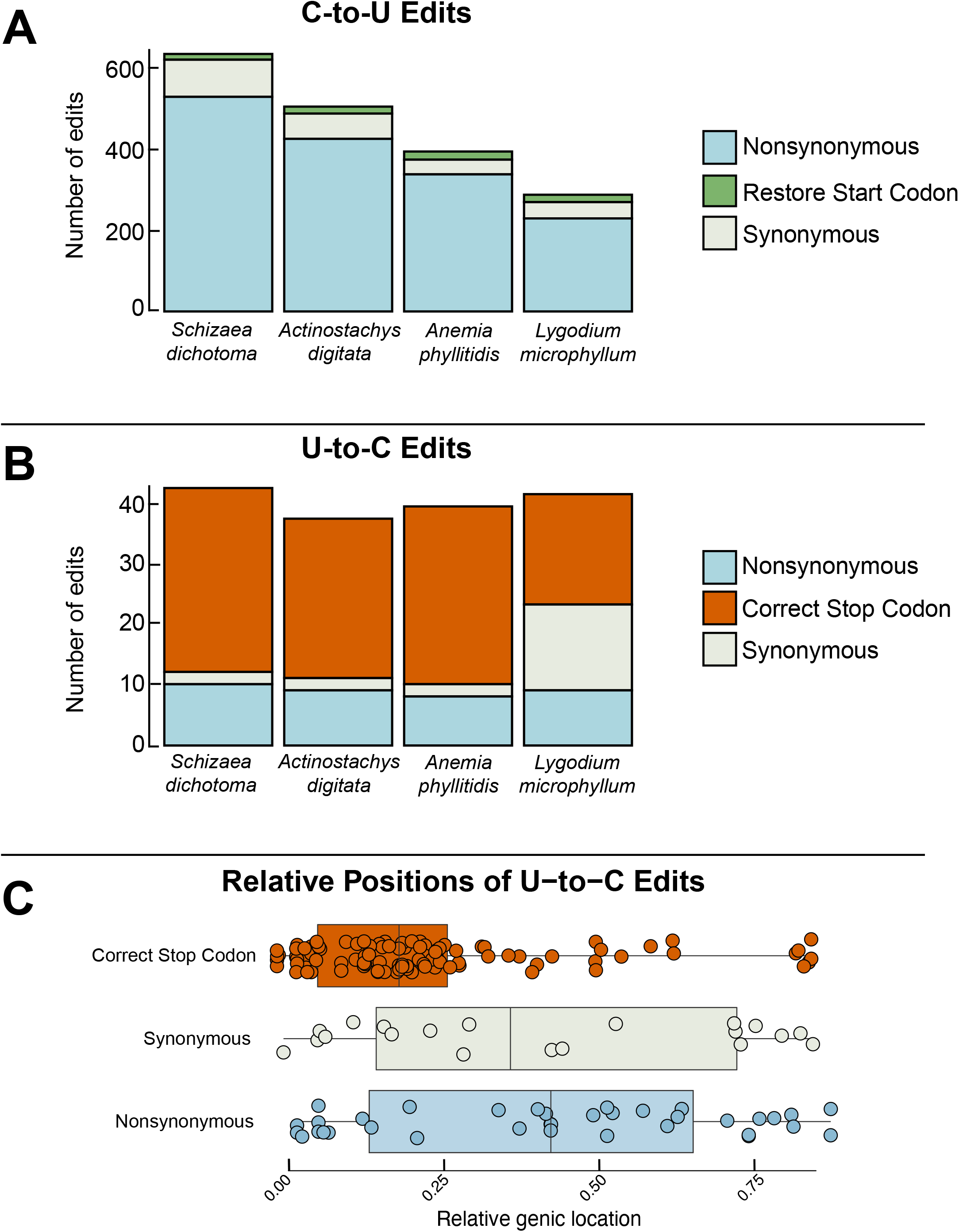
Proportion of RNA editing sites resulting in nonsynonymous (light blue), synonymous (grey) changes, as well as cryptic start codon restoration (green, C-to-U only) and internal stop codon correction (orange; U-to-C only). A shows C-to-U edits, B shows U-to-C edits. C shows the relative genic positions of U-to-C edits grouped by codon change type.

RNA editing efficiencies—defined as the percentage of mapped RNA reads at a given site that are edited—were generally lower in the Schizaeaceae compared to *Lygodium* and *Anemia*, particularly for U-to-C edits (Fig. 5). Across all species, synonymous edits were typically edited at low efficiency regardless of direction. Nonsynonymous C-to-U edits occurred at slightly higher efficiencies in *L. microphyllum* and *A. phyllitidis* than edits that restore cryptic start codons, whereas in *S. dichotoma* and *A. digitata*, their efficiencies were more comparable. U-to-C edits in the Schizaeaceae were often edited at less than 50% efficiency, contrasting with the higher efficiencies observed in *A. phyllitidis* and *L. microphyllum* (Fig. 5).

**Figure 5.**
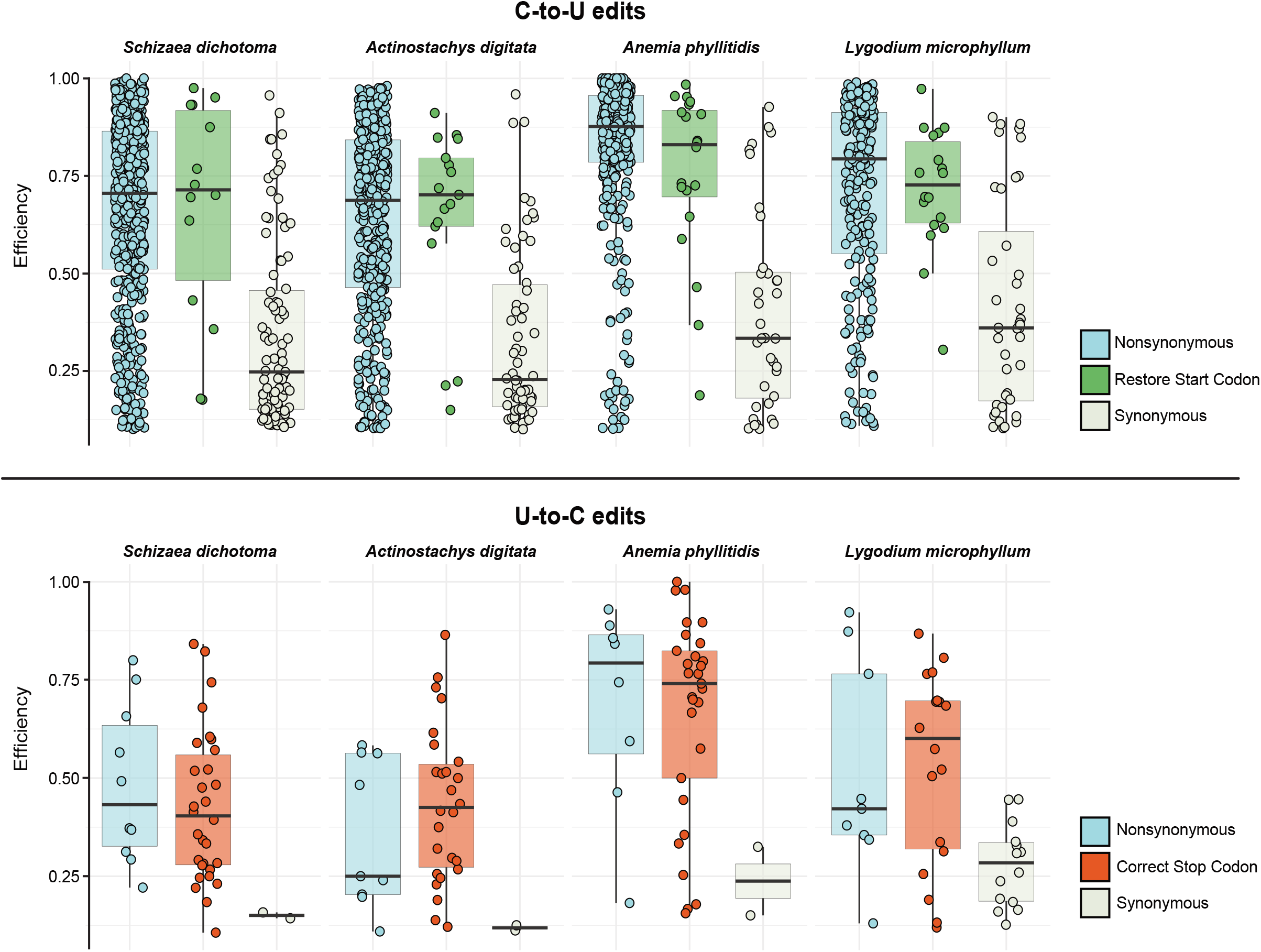
RNA efficiencies for four Schizaeales species. The top panel shows C-to-U editing efficiencies with nonsynonymous edits in light blue, synonymous edits in grey, and edits that restore start codons in green. The bottom panel shows U-to-C editing efficiencies with nonsynonymous edits in light blue, synonymous edits in grey, and edits that correct internal stop codons in orange.

Using gene alignments, we first compared nucleotide diversity (π) before and after editing and found that π from DNA alignments was consistently higher than π from post-edited RNA alignments for nearly all genes and across all species pairs (Fig. 6). We then assessed the conservation of individual edits and discovered that most nonsynonymous C-to-U and U-to-C edits were lineage specific (Fig. 7). 40 genes across Schizaeales require C-to-U editing to restore the start codon in at least one species. Each species individually possesses between 20 and 14 and start codon edits (Fig. 7). These edits appeared slightly more conserved, with five start codon edits found in all species except *L. microphyllum* (Fig. 7). We found no start codon edits to be conserved across all species. We recovered a total of 49 unique internal stop codons requiring correction by U-to-C editing across the Schizaeales, ranging from 31 in *S. dichotoma* to 19 in *L. microphyllum* (Fig. 7). Of these, 7 are conserved across the order, with another 7 shared exclusively by the Schizaeaceae, and 5 more conserved between Schizaeaceae and *A. phyllitidis* (Fig. 7).

**Figure 6.**
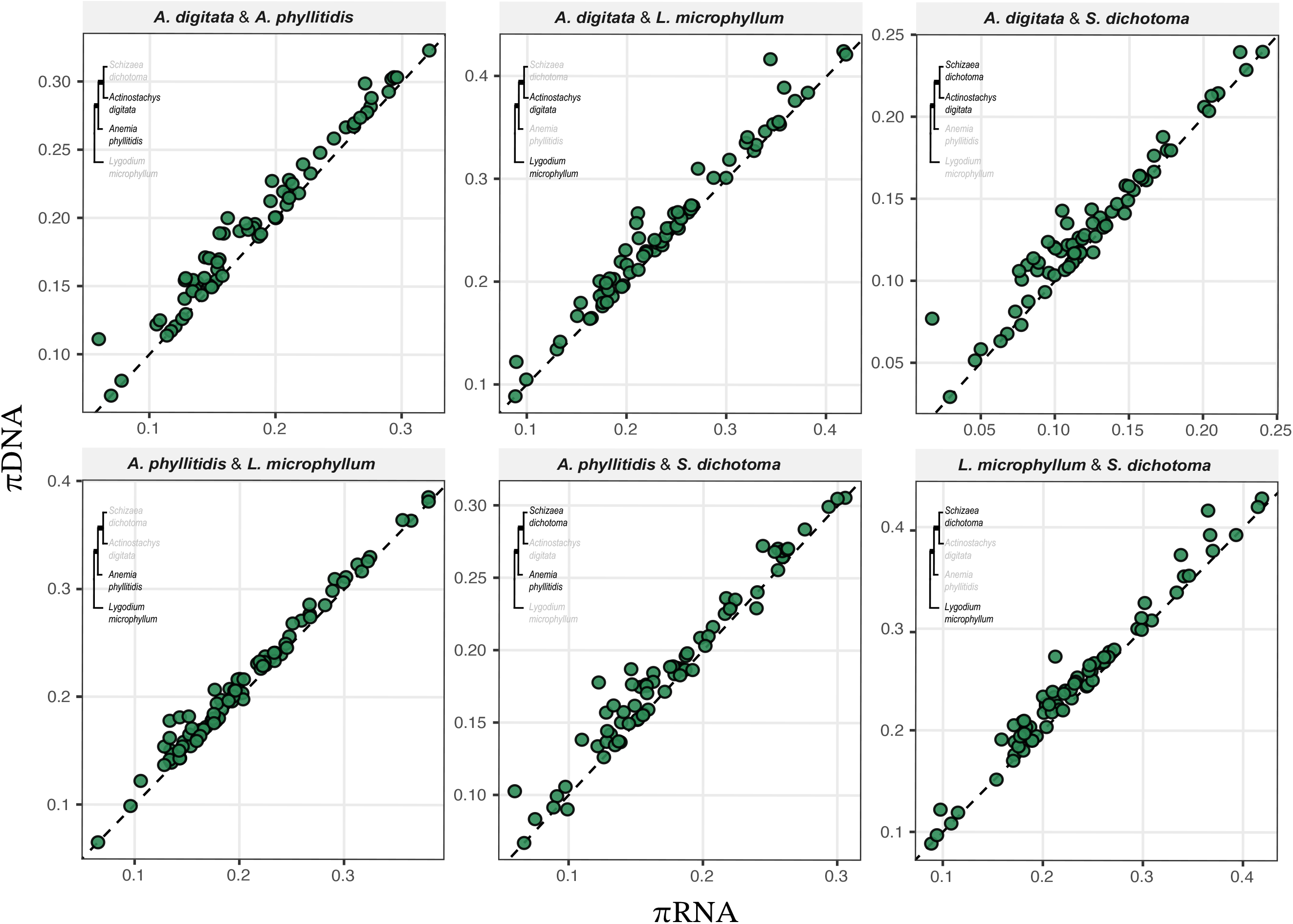
Pairwise nucleotide diversity (π) calculated on DNA alignments and post-editing RNA alignments across six pairs of Schizaeales species for each plastid gene. DNA π is shown on the y-axis and RNA π on the x-axis. The dashed trendline represents the 1:1 line. Points above this line represent genes with a higher DNA π than RNA π and points below represent genes with a lower DNA π than RNA π.

**Figure 7.**
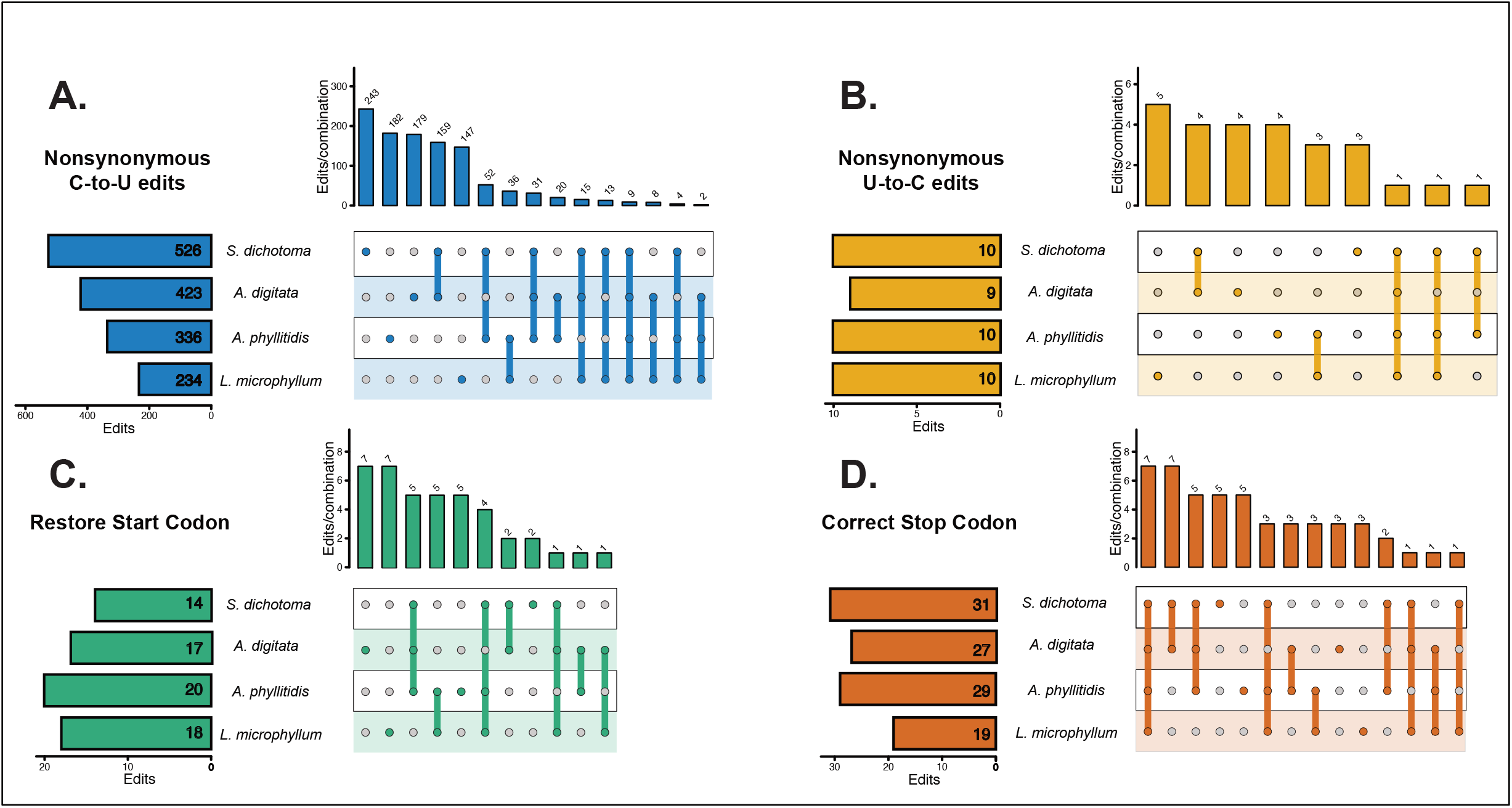
Proportion of shared RNA editing sites across Schizaeales. Nonsynonymous C-to-U edits are shown in blue (A), nonsynonymous U-to-C edits in yellow (B), C-to-U edits at start codons in green (C), and U-to-C edits at internal stop codons in dark orange (D). Horizontal bars to the left of species abbreviations show the number of edits present in each species. Vertical bars denote how many edits are shared by an exclusive group of taxa, defined by the connected dots.

Using a dataset with expanded sampling to include additional *Schizaea, Actinostachys*, and *Lygodium* species, we found statistically significant evidence for relaxed selection in the Schizaeaceae for 28 genes (Fig. 8). Support for the relaxation of selection in Schizaeaceae was found in genes for nearly every functional group but was particularly pronounced for photosystem genes (*psa* and *psb*) as well as ATP synthase genes (*atp*; Fig. 8). Plastid-encoded RNA polymerase genes (*rpo*) and *ycf1* exhibited some of the highest pairwise nonsynonymous C-to-U editing differences between Schizaeaceae and non-Schizaeaceae species, however these are also among the longest genes in chloroplast genomes. To account for gene length, we ran a multiple linear regression to test the relationship between the relaxation parameter (*k*) and the pairwise editing difference between Schizaeaceae and non-Schizaeaceae species in genes under relaxed selection, using gene length as a covariate. *k* represents the strength of selection relative to a reference set of branches—here the *Lygodium* and *Anemia* branches—and for genes where selection is relaxed, it ranges from 0 to approaching 1. *k* = 0 represents a complete relaxation of selection whereas values close to 1 represent genes where selection is only slightly relaxed. For the relaxed genes, *k* varies between 0 in some genes to 0.78. For 3 out of 4 comparisons, we find a significant negative relationship between *k* and the pairwise editing difference (Fig. 9). The only comparison that did not meet a p < 0.05 significance threshold was *S. dichotoma* vs *A. phyllitidis*, however the p-value here is quite close at 0.0763 (Fig. 9).

**Figure 8.**
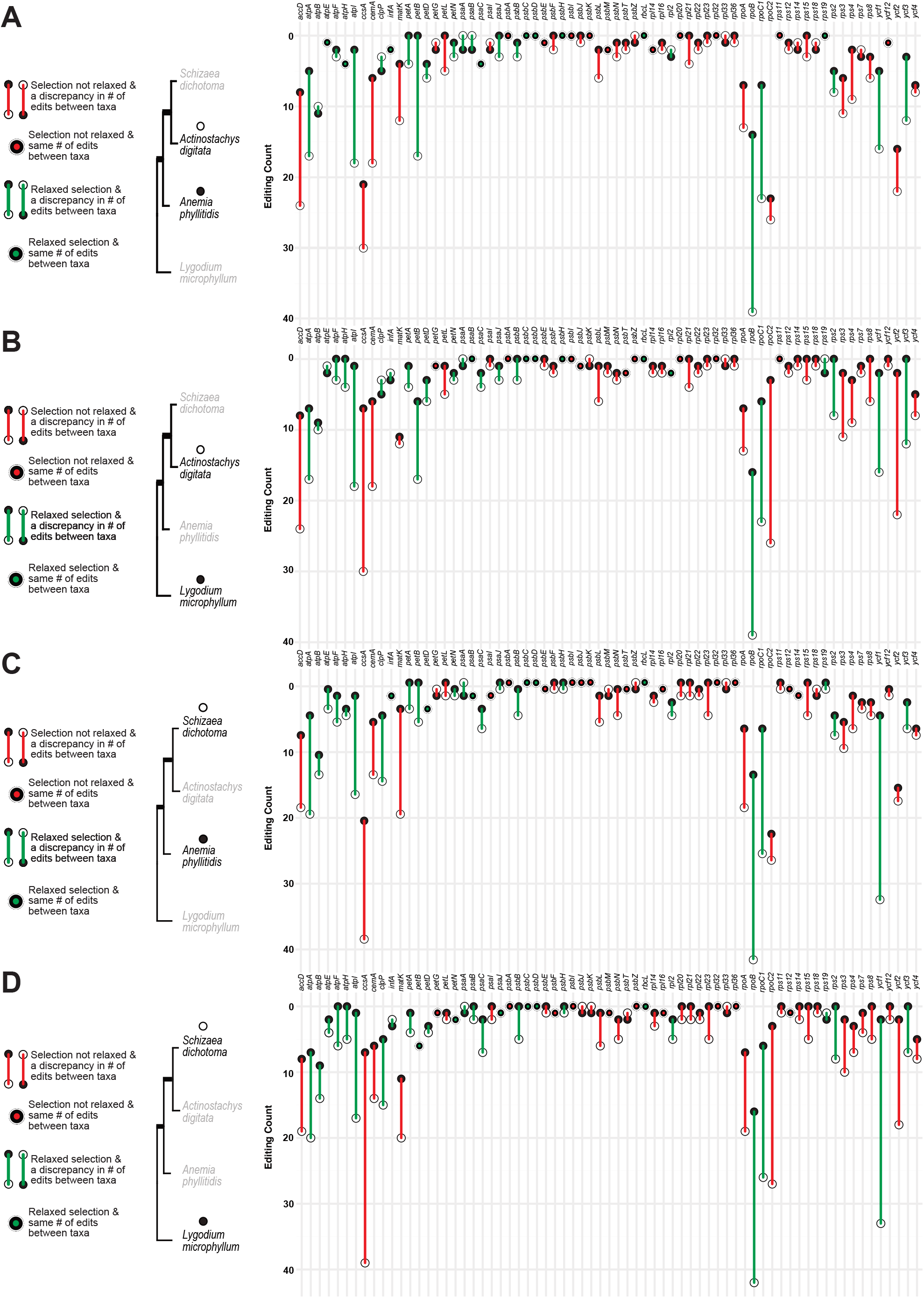
Pairwise differences in the number of C-to-U edits for each gene for every Schizaeaceae vs. non-Schizaeaceae comarison. Open circles are used for Schizaeaceae species (*Schizaea dichotoma* or *Actinostachys digitata*) and dark circles for non-Schizaeaceae species (*Anemia phyllitidis* or *Lygodium microphyllum*). Genes under relaxed selection in the Schizaeaceae are shown in green while genes not under relaxed selection are in red.

**Figure 9.**
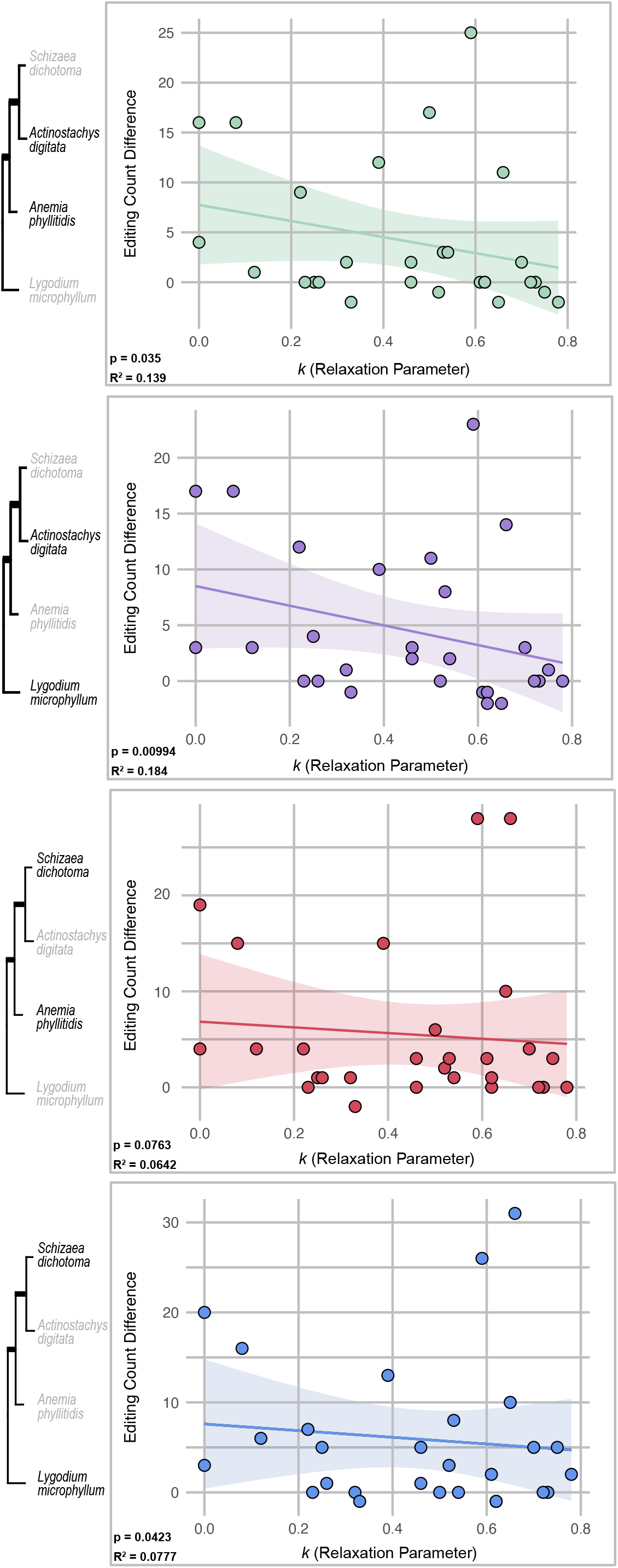
Relationship between relaxation parameter k and the pairwise difference in the number of C-to-U edits in every gene under relaxed selection. All Schizaeaceae vs. non-Schizaeaceae comparisons are shown.

For genes that are consistently either in single-copy regions or the IR across Schizaeales, we found the mean relative MRCA distances to generally be lower in *Lygodium*, than *Actinostachys* and *Schizaea* (Fig. 10A,B). However, for the five genes that have been translocated into the IR in *Schizaea* specifically, *Schizaea* typically has a much lower mean relative MRCA distance than *Actinostachys* or *Lygodium*, with *psaC* being the only exception (Fig. 10C). Three genes were translocated into the IR for both *Actinostachys* and *Schizaea*. Here *Actinostachys* consistently has a lower mean relative MRCA distance than *Schizaea* and *Lygodium* (Fig. 10D). For *matK, Schizaea* has a lower mean relative MRCA distance than *Lygodium*, very close to that of *Actinostachys*, but it has the highest mean relative MRCA distance for *rpl32* and *rpl21* (Fig. 10D). For translocated genes, C-to-U RNA editing levels are consistently higher in species where that gene resides in the IR as opposed to the LSC or SSC (Fig. 10E).

**Figure 10.**
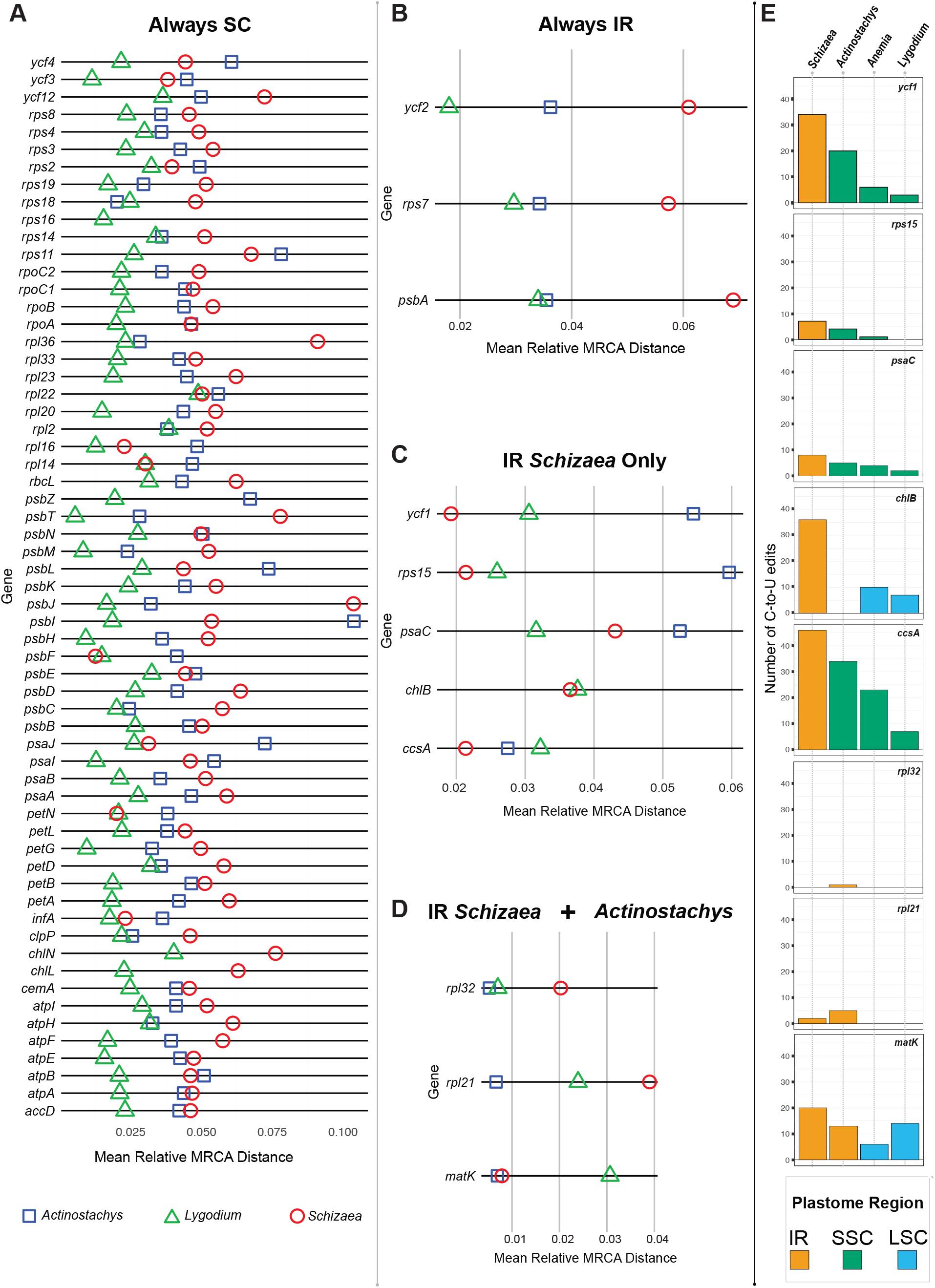
Panels A–D show the mean relative distance to the most recent common ancestor (MRCA) for *Lygodium* (green triangle), *Actinostachys* (blue square), and *Schizaea* (red circle) for each plastid gene shared among the three genera. Panel A shows genes that are consistently found in the single copy regions, panel B shows genes found consistently in the inverted repeat (IR), C shows genes translocated into the IR in *Schizaea* only, and D shows genes translocated into the IR in Schizaea and *Actinostachys*. Panel E shows the amount of RNA editing for every gene that has been translocated for four Schizaeales species, colored by what region that gene resides in each species. IR is colored in gold, the small single copy region (SSC) in green, and the large single copy region (LSC) in blue.

### Schizaeales PPR protein diversity

Using *de novo* assembled transcriptomes, we identified PPR genes in each of the four Schizaeales species. Using this method, we identified thousands of PLS class PPR proteins in each species (Table 1). However, we found very few of these to contain catalytic editing domains (DYW for C-to-U editing and DYW:KP for U-to-C editing; Table 1). At most we found 35 full-length C-to-U editing factors in *L. microphyllum* to as few as 4 in *S. dichotoma* (Table 1). Similarly, we found at most 16 full-length U-to-C editing factors in *L. microphyllum* to as few as 4 in *A. digitata* (Table 1).

**Table 1:** Schizaeaceae PPR proteins. PLS sequences are PPR proteins with PLS triplets, with a C-terminal truncation with no distinctive features of C-to-U or U-to-C editing proteins. The E+ sequences are PPR proteins with PLS triplets, with a C-terminal truncation ending in an E motif with signatures indicative of C-to-U editing proteins. The E+:KP sequences are PPR proteins with PLS triplets, with a C-terminal truncation ending in an E motif with signatures indicative of U-to-C editing proteins. The DYW sequences are PPR proteins with a DYW domain. The DYW:KP sequences are PPR proteins with a DYW:KP domain.

|  | PLS | E+ | E+:KP | DYW | DYW:KP | Total |
| --- | --- | --- | --- | --- | --- | --- |
| *Schizaea dichotoma* | 2304 | 18 | 16 | 4 | 6 | 2348 |
| *Actinostachys digitata* | 2182 | 13 | 16 | 6 | 4 | 2221 |
| *Anemia phyllitidis* | 3095 | 32 | 34 | 13 | 12 | 3186 |
| *Lygodium microphyllum* | 1266 | 280 | 214 | 35 | 16 | 1811 |

## Discussion

Over the past three decades, substantial progress has advanced our understanding of the evolutionary dynamics of RNA editing in plant organellar genomes (Small et al. 2020). A prevailing model proposes that editing sites originate through constructive neutral evolution (CNE), a ratchet-like process driven by neutral mutations; (Covello and Gray 1993). However, the overall abundance of editing sites in a plastome is shaped by more than just their mode of origin. The high frequency of C-to-T mutations in AT-rich plastid genomes (Huang et al. 2005) continuously erodes editing sites by converting editable cytidines into thymidines, thereby eliminating the need for editing. At the same time, not all potential editing sites are tolerated—selection may quickly purge sites that disrupt gene expression or impose burdens on highly transcribed genes. Thus, the observed distribution of RNA editing sites emerges from three competing forces: neutral origination via CNE, purifying selection that eliminates deleterious edits, and backmutation (e.g., C-to-T substitutions) that progressively erodes established sites (Mower 2008; Ishibashi et al. 2019; Fauskee et al. 2025).

Despite these advances, plastid RNA editing evolution is well characterized in fully photosynthetic lineages but poorly understood in species with reduced or absent photosynthetic capacity. In these systems, plastid genomes often experience gene loss, and the selective constraints on retained plastid genes—particularly photosynthesis-related genes—are presumably relaxed.This raises a key question: under relaxed selection, do editing sites accumulate more readily via CNE, or do gene losses and elevated mutation rates lead to a net reduction in editing sites? Addressing these questions is essential for understanding RNA editing evolution in functionally reduced plastid genomes.

Ferns are an excellent model for studying plastid RNA editing evolution. They retain higher C-to-U editing levels than seed plants and are among the few vascular plant lineages that retain U-to-C editing, which has been lost in seed plants. We investigated plastid RNA editing in the fern order Schizaeales, which comprises three families: Schizaeaceae, Anemiaceae, and Lygodiaceae (PPG I 2016). While all species in Anemiaceae and Lygodiaceae exhibit typical photosynthesis in both gametophytes and sporophytes, Schizaeaceae includes several lineages with achlorophyllous, subterranean, mycoheterotrophic gametophytes (Bierhorst 1975; reviewed in Ke et al. 2022). Here we asked whether RNA editing levels differ between fully autotrophic and partially heterotrophic ferns, and what evolutionary forces might drive those differences.

Our plastome assemblies and annotations corroborate findings initially reported by Labiak and Karol (2017) as well as by Ke et al. (2022). Specifically, we recovered the same IR expansion and plastome reduction in *Actinostachys digitata* as well as the more extreme IR expansion and dramatic SSC reduction in *Schizaea dichotoma* (Fig. 1). We also recovered the same gene losses in the two Schizaeaceae species as previously reported, including the entire loss of the electron transport (*ndh*) gene suite, *psaM, rps16*, and *ycf66*, along with the specific loss of the three chlorophyll biosynthesis (*chl*) genes in *A. digitata* (Fig. 2). Despite these extensive gene losses across Schizaeaceae, both species showed a dramatic increase in C-to-U RNA editing levels. The *S. dichotoma* plastome harbored more than double the number of C-to-U edits found in *L. microphyllum*, despite encoding 14 fewer protein-coding genes (Fig. 3). In contrast, there was strikingly little variation in the number of U-to-C edits across all species examined (Fig. 3), supporting the hypothesis that C-to-U and U-to-C editing are likely shaped by different evolutionary pressures (Fauskee et al. 2021,2025).

Consistent with other fern lineages (Fauskee et al. 2025), most C-to-U edits caused nonsynonymous amino acid substitutions, with some restoring start codons, whereas U-to-C edits primarily corrected internal stop codons (Fig. 4). Both Schizaeaceae species exhibited lower C-to-U RNA editing efficiencies than Anemia and Lygodium, across all edit types: nonsynonymous edits, synonymous edits, and edits restoring start codons (Fig. 5). Schizaeaceae also exhibited an even more dramatic reduction in U-to-C editing efficiency relative to Anemia and Lygodium (Fig. 5). Here most U-to-C edits are edited with below 50% efficiency for Schizaeaceae, whereas many U-to-C edits in Anemia and Lygodium have much higher efficiencies (Fig. 5). Across the Schizaeales, RNA editing efficiencies at internal stop codons are quite variable, biased toward the 5’ end of the transcript, and appear to be evolutionarily conserved (Figs. 4,5). These observations are consistent with the hypothesis that selective U-to-C editing of internal stop codons functions in plastid gene regulation in ferns (Fauskee et al. 2025; Kwok van der Giezen et al. 2025).

Schizaeaceae exhibited elevated C-to-U RNA editing site numbers relative to Anemia and Lygodium (Fig. 3), but reduced editing efficiencies (Fig. 5). We asked whether the increased number of edits reflects a greater accumulation and retention of true editing sites (through higher rates of site gain and/or lower rates of site loss), or “promiscuous” off-target editing by PPR proteins at mismatched binding sites. Off-target edits would likely show low efficiency because of imperfect binding site matches. To assess these alternatives, we calculated pairwise nucleotide diversity (π) for each gene and compared π calculated from DNA alignments to π from post-edited RNA alignments (Fig. 6). If the increased editing reflects off-target activity, then the post-edited RNA should show higher π than the genomic DNA, as these non-corrective edits would introduce additional sequence variation in the mRNA relative to the DNA. Alternatively, if the higher edit numbers stem from a higher rate of gaining edits through CNE and/or a decreased rate of losing edits through backmutation, we would expect to see higher DNA π than RNA π. This is because edits acquired through CNE are predominantly corrective so differences in the DNA will be resolved in the RNA via RNA editing. Across nearly all genes and pairwise comparisons, DNA π exceedsed RNA π (Fig. 6), indicating limited off-target editing. Instead, the difference in the amount of RNA editing is likely due to some combination of increased rates of editing site establishment through CNE and decreased rate of losing edits through backmutation. Currently, we lack the sampling to distinguish between the two possibilities, though it is likely both play a significant role in the overall editing distribution.

The overwhelming majority of plastid RNA editing events in Schizaeales are nonsynonymous C-to-U edits (Fig. 4). These edits are not evolutionarily conserved—each species harbors over 100 unique editing sites not shared with any other species (Fig. 7). Our selection analyses show that 28 genes are under relaxed selection in Schizaeaceae relative to Anemia and Lygodium (Fig. 8). Many of the largest pairwise differences in the number of nonsynonymous C-to-U edits—comparing Schizaeaceae to Anemia and Lygodium —occur in these relaxed genes, particularly ycf1, rpoB, and rpoC1 (Fig. 8). Furthermore, in all but one pairwise comparison (Schizaea vs. Anemia), we find a statistically significant negative relationship between editing count differences and k, suggesting that genes under relaxed selection accumulated more nonsynonymous C-to-U edits in Schizaeaceae than in their photosynthetic relatives (Fig. 9).

This result is somewhat unexpected. Previous studies in both ferns (Fauskee et al. 2025) and angiosperms (Mower 2008; Ishibashi et al. 2019) report a consistent trend of C-to-U editing site loss over time, with faster-evolving lineages typically harboring fewer such edits. The elevated substitution rates in Schizaeaceae (Fig. 3), suggest accelerated backmutation, which would be expected to reduce editing levels. Instead, we observe the opposite: Schizaeaceae species possess a higher number of C-to-U edits. This pattern suggests that relaxed selection facilitates the establishment of new editing sites. Supporting this interpretation, genes under weaker selection tend to accumulate more edits in Schizaeaceae compared to their orthologs in Anemia and Lygodium (Fig. 9) and the DNA-RNA π comparison confirms that the editing difference is indeed due to novel unique edits, not off-target editing (Fig. 6). Taken together, these results point to a balance between selection and the neutral origin of edits via constructive neutral evolution (CNE). When selection is relaxed, newly arising editing sites are more likely to persist, and their accumulation scales with the degree of relaxation (Fig. 9). Moreover, PLS-class PPR proteins are known to strongly prefer binding to uridines (Dennis et al. in review). The high rate of C-to-T substitutions in plastid genomes—especially in genes under relaxed selection—may generate additional U-rich motifs that PPR proteins can recognize, thereby expanding the pool of potential editing sites. Thus, the elevated number of C-to-U edits in Schizaeaceae likely reflects an accelerated CNE ratchet, wherein relaxed selection simultaneously increases mutation rate, increases the availability of editable U-rich motifs, and permits the persistence of editing sites that would be eliminated under stronger constraint. In other words, relaxed selection both increases the number of potential PPR binding sitesthrough C-to-T substitutions that generate U-rich motifs —and reduces barriers to retaining established edits, together promoting the extraordinary accumulation of editing sites in Schizaeaceae.

Beyond relaxed selection, a second factor contributing to elevated RNA editing levels in Schizaeaceae is expansion of the inverted repeat (IR). Using our expanded sampling, we calculated the mean branch length from each Lygodium, Actinostachys, and Schizaea species to the most recent common ancestor (MRCA) of their respective genus. For genes consistently located in either the IR or the single-copy regions throughout their evolution, Schizaea and Actinostachys tend to have relatively longer branches than Lygodium (Fig. 10 A,B). However, genes translocated into the IR showed the opposite pattern. In both Schizaea and Actinostachys, IR-relocated genes generally showed shorter relative mean branch lengths than the same genes in Lygodium (Fig. 10C,D), suggesting reduced substitution rates following IR integration. This agrees with previous findings that IR-relocated genes experience reduced substitution rates in ferns (Li et al. 2016). Consequently, for every case of IR translocation, C-to-U editing levels were consistently higher in the species harboring the gene in the IR (Fig. 10E). These results suggest that the reduced substitution rates associated with IR translocation slows the erosion of C-to-U editing sites. In effect, the IR “protects” editing sites from being lost over time, because the lower mutation rate reduces the chance of backmutation at edited sites.

Interestingly, a different pattern is observed in the lycophyte Phylloglossum drummondii, which has a partially subterranean and tuberous gametophye stage. In contrast with the Schizaeaceae, P. drummondii retains most of the conserved plastid genes, including the often-lost ndh suite, and exhibits relatively low levels of RNA editing (Kwok van der Giezen et al. 2024). Although interpretation is complicated by a generally low number of plastid RNA editing sites in huperzioid lycophytes overall, P. drummondii provides a notable exception whereby the transition to a partially non-photosynthetic lifestyle is not accompanied by plastid genome degradation or elevated RNA editing.

A second unexpected finding was the striking lack of full-length PPR RNA editing factors in Schizaeales (Table 1). Across Schizaeales, we recovered far fewer PPR proteins with C-terminal editing domains than would be expected given the number of plastid RNA editing sites. This shortfall is even greater when taking into account the extensive RNA editing known to occur in the mitochondrial genome (Knie et al. 2016; Guo et al. 2017; Zumkeller et al. 2023). However, we did recover a large number of PLS-class PPR proteins lacking their catalytic C-terminal domains which could account for the high number of editing sites found in both the plastid and presumably mitochondrial genomes. These PLS proteins contain the RNA-binding arrays responsible for recognizing editing sites by binding to upstream cis sequences. This strongly suggests that in Schizaeales—and likely in ferns more broadly (Li et al. 2018; Gutmann et al. 2020)—the RNA-binding and catalytic functions of RNA editing are carried out by separate proteins assembled in trans. In the distantly related Salviniales ferns, the number of DYW and DYW:KP domains responsible for C-to-U and U-to-C conversion, respectively, are far outnumbered by the abundance of RNA editing sites and PLS PPR proteins (Li et al. 2018). This mismatch in Schizaeales is even more striking, where as few as 4 DYW domains appear to serve 631 C-to-U editing sites in the Schizaea dichotoma chloroplast and mitochondrial transcriptome. In angiosperms, PPR proteins interact with several auxiliary factors such as RIP/MORF, ORRM and OZ1 proteins. However, these are absent in seed-free plants where PLS motif tracts tend to be longer and may act independently. Some seed-free lineages, such as the hornwort Anthoceros, have a near one-to-one ratio of full-length U-to-C editing factors, but a mismatch in the number of DYW domains to C-to-U editing sites (Gutmann et al. 2020). We cannot exclude the possibility of a seed-free plant-specific RNA editing co-factor protein or protein complex, but it seems apparent that DYW and DYW:KP domains can act as ‘donor’ catalytic domains for C-to-U and U-to-C editing. Schizaeales ferns present a good model for studying the protein-protein interface between PLS PPR proteins and donor DYW/DYW:KP domains. Moreover, decoupling the PLS binding domain from a fixed catalytic identity (e.g., C-to-U vs. U-to-C) could allow for greater flexibility in site targeting and may facilitate the rapid accumulation of RNA editing sites. This modularity may help explain why fern plastomes harbor far more editing sites than those of angiosperms.

## Conclusions

Together, our findings provide new insight into the evolutionary dynamics that shape plastid RNA editing in ferns and, more broadly, in land plants. By examining species with non-photosynthetic gametophytes, we reveal how relaxed selection can shift the balance between the neutral accumulation of editing sites and the forces that typically limit their presence. Previous studies have emphasized the gradual erosion of editing sites over time, primarily driven by high C-to-T mutation rates that eliminate sites through backmutation. In addition, selection can prevent the establishment of editing sites in the first place, particularly in highly-expressed genes where the additional complexity of RNA editing may be less tolerable. Our results demonstrate that when selection is relaxed, as in many plastid genes of Schizaeaceae, this mutational erosion can be outpaced by the neutral gain of editing sites through constructive neutral evolution (CNE). In these lineages, relaxed selective constraints enable the CNE ratchet to turn more freely, enabling newly arising editing sites to establish and persist. Moreover, PPR RNA editing factors preferentially bind uridine, and the higher rate of C-to-T substitutions under relaxed selection may further expand the number of potential cis–binding domains that these factors can recognize, further accelerating the accumulation of plastid RNA edits in Schizaeaceae. As a result, we observe significantly more C-to-U editing events in Schizaeaceae species, even though they possess fewer protein-coding genes and have higher overall substitution rates than their fully-photosynthetic relatives. We also show that structural changes to the plastome—specifically, expansion of the inverted repeat (IR)—contribute to elevated editing levels by decelerating substitution rates. In Schizaea and Actinostachys, genes translocated into the IR consistently exhibit shorter branch lengths relative to Lygodium than their single-copy counterparts, indicating a deceleration of molecular evolution. This substitution rate slowdown likely delays the progressive loss of editing sites by reducing the frequency of backmutation at edited cytidines. In effect, IR expansion may “preserve” editing sites present at the time of translocation by slowing the mutational clock. Finally, we find that Schizaeales plastomes contain far more editing sites than can be explained by the number of full-length PPR editing factors alone. Instead, these species harbor many PLS-class PPR proteins that lack catalytic domains, suggesting that RNA-binding and catalytic functions are modular and assembled in trans. This arrangement may provide greater flexibility in targeting RNA editing sites and possibly allow a single catalytic component to serve multiple editing complexes, potentially facilitating the rapid accumulation and retention of editing sites. Altogether, our results highlight how both relaxed selection and plastome structural arrangements can enable the expansion of plastid editomes, and how examining partially non-photosynthetic systems helps to illuminate the interplay between neutrality, selection, and molecular machinery in shaping RNA editing evolution.

